# Acute and chronic toxicity assessment of benzylpenicillin G residues in heat-treated animal food products

**DOI:** 10.1101/191346

## Abstract

The current level of penicillin use and its persisting residues in livestock is potentially concerning; the toxicity of penicillin residues in heat-treated animal food products (HAFP) is yet to be elucidated. In this study, the acute and chronic toxicity of benzylpenicillin G (BPG) residues in HAFP was investigated in a mouse model. The calculated LD_50_ of BPG heated to cooking temperature (BPHCT) was 933.04 mg kg^-1^ [b.w.] intraperitoneally corresponding to 3.75 times lower than its prototype. Mice fed on the experimental diet containing heat-treated beef with high BPG levels for 6 months displayed a reduction in body weight and altered serum values indicating for liver and renal function. Further, the organ ratios of intestinal and spleen were increased. Histopathological changes were observed in the liver, lung and parenchyma testis tissue. BPHCT residue induced sperm aberration and micronucleated polychromatic erythrocytes formation. Present results indicate that prolonged exposure of BPHCT at higher levels of residue might have an impact on public health. Importantly the toxic concentrations of BPHCT are relatively high compared with levels that would result from the degradation of antibiotic residues in meat from animals that have received a therapeutic dose of BPG.

## 1. Introduction

The effect of the veterinary drug residues on human health is of increasing concern due to the growing consumption of animal derived products (Baynes et al., 2016). Penicillin, a β-lactam antibiotic, has been widely used in food-producing animals (cattle, pigs and poultry) as a veterinary antibacterial agent to control diseases due to its high bacterial killing efficiency, relatively low toxicity and cost (Edwards and Brownlee, 1946; Dunlop et al., 1998). Penicillins also work as growth promoters to enhance the feed efficiency in animal husbandry (Barton, 2000).

With the development of drug-resistance in bacteria, penicillin is often used at higher dosage rates than those indicated on the manufacture recommendations (Chiesa et al., 2006). The typical dose of penicillin used in cattle by intramuscular injection is approximately 3.5 to 10 times greater than the US approved dose (Payne et al., 2006). It was reported that during the years of 2005-2009, nine countries in the EU consumed a total of 11,342 tonnes antibiotics, of which 18.96% was penicillin, with the majority used in Finland, Netherlands and the United Kingdom (Grave et al., 2012). In 2011, total sales and distribution of penicillin approved for use in food-producing animals was 885 tonnes in the USA (FDA, 2014). In China, 4960 tonnes of penicillin was used in animals during the year of 2013 (Zhang et al., 2015).

With the large amount of penicillin used, there are increased risks of exceeding the Maximum Residue Limit (MRL) range (4 µg kg^-1^ and 50 µg kg^-1^ in milk and muscle, respectively (Di Corcia and Nazzari, 2002; Commission, 2009)). According to the USDA National Residue Program report, penicillin residues in excess of legal permitted levels were detected in 22% of the total number of violations in the US (Li et al., 2017a). One study reported that in milk, 28% of collected samples were antibiotic positive, of which 11% were considered non-compliant with current European Union regulations of penicillin (Junza et al., 2014). Becker et al reported that the highest concentration of penicillin residues in bovine kidney and milk were 1200 µg kg^-1^ and 536 µg kg^-1^, respectively (Becker et al., 2004). In Italy, Ghidini et al. found that the highest concentration of penicillin residues in milk was 6240 µg L^-1^ with the mean concentration of 510.2 µg L^-1^ of collected samples (Ghidini et al., 2003). Samanidou et al. reported that the concentrations of penicillin residues in beef were 156 µg kg^-1^ and 489 µg kg^-1^, which were collected in Greek local markets (Samanidou et al., 2007). Myllyniemi et al. reported that the number of penicillin residues in kidney and muscle samples above 400 µg kg^-1^ were 10 and 3, respectively, which were from slaughterhouses around Finland (Myllyniemi et al., 2000). Penicillin residues have been reported in chicken from ‘organic’ farms with a concentration of 1.3 ng g^-1^, which was labelled as antibiotic-free in Hong Kong local markets (Li et al., 2017b). In some developing countries, the level of penicillin residues in animal derived products may be under reported due to the lack of quality assurance programs (Kabir et al., 2004; Kang’ethe et al., 2005; Babapour et al., 2012). In addition, β-lactams antibiotics were detected in 10% of the urine samples from children aged 8-9 in China, with the highest concentration of 40000 ng mL^-1^ for ampicillin (Wang et al., 2015b).

Penicillin is unstable, with many studies highlighting the heat instability of penicillin in animal derived products. Ludger et al. found that residues from penicillin, penillic, penicilloic and penilloic acids formed in the milk and yoghurt after heat treatment and fermentation by LC-MS/MS detection method (Grunwald and Petz, 2003). Rose et al. suggested that temperature higher than 65 □ would affect the structure of penicillin, making its half life varying from 15 to 60 min leading to generation to penicilloic acid as the major breakdown product in cooked food (Rose et al., 1997). Other studies have reported additional adverse effects of penicillin residues in animal derived products. It has been reported that penicillin residues in food could cause allergic reaction, which is known to be mediated by IgE antibody and the soluble factor like IL-4 or IL-13 (Pene et al., 1988; Dayan, 1993; Punnonen et al., 1993; Pawankar et al., 1997; Guéant-Rodriguez et al., 2006). Mund et al. reported that some severe clinical symptoms such as dermatitis, cutaneous eruptions, anaphylaxis and gastro-intestinal symptoms in humans were the results of penicillin residues in poultry products (Mund et al., 2016). Aarestrup et al. emphazised that penicillin residues could also facilitate the spread of antibiotic resistance genes from animals to humans through food chain (Aarestrup et al., 2008). To our knowledge, there is only one study that reported the toxicity of one major degradation product of BPHCT-benzylpenicilloic acid, which showed toxicity *in vivo* and *in vitro* (Cui et al., 2018).

In this study, we investigated both acute and chronic toxicity studies in mice to understand the possible adverse effect posed by BPHCT to human health.

## 2. Materials and Methods

### 2.1 Chemicals and reagents

Benzylpenicillin G (BPG, molecular weight: 356.37; 1.6 million units 0.96 g^-1^; CAS No.: 61-33-6; purity >99%) was purchased from North China Pharmaceutical Co., Ltd and stored in a dark and dry place at 4□ for subsequent tests. Phosphate buffered solution (PBS, Cat. No. PYG0021), poly-lysine treated slides (Cat. No. AR1065), mouse interleukin-4 (IL-4, Cat. No. EK0405) and interleukin-13 (IL-13, Cat. No. EK0425) ELISA Kits were purchased from BOSTER Biological Technology Co. Ltd. China. Mouse immunoglobulin E (IgE, Cat. No. 88-50460-22) ELISA kit was purchased from Thermo Fisher Scientific Co. Ltd, USA.

### 2.2 Preparation of test substance

BPG (1.6 million units 0.96 g^-1^) was dissolved in 2 mL of 0.9% saline solution and used for the study. Normal diet containing BPHCT and beef were served as the experimental diets and prepared as follows. Briefly, BPG (480 mg mL^-1^) was injected into raw beef (1 g kg^-1^ [bw] day^-1^) at different dose levels and boiled with beef for 20 min. The details of dose of BPG and weight of beef used in the study are shown in the supplementary material. The heat-treated mixtures together with the boiling water were all added into the normal diet, mixed evenly and reshaped to form the experimental dry diets. Normal diet mixed with heat-treated beef and 0.9% saline solution was employed as the negative control.

### 2.3 Animals

ICR mice were obtained from Laboratory Animal Centre of Jilin University. The animals were 23-25 g in weight and clinically examined to rule out any ailment and only ones without any signs of diseases were employed in the further study. Mice were housed at 22 □, 50%-70% humidity and artificial lighting was on 12 h day and night cycle. Mice were fed with the normal diet and unlimited water. An adaptation period of at least 7 days was employed before the start of actual dosing. The animal experiments studies were approved by the Animal Care and Use Committee of Changchun Experimental Animal Centre, Jilin University. The experiments were designed and conducted in compliance with the OECD Principles and Good Laboratory Practice (GLP).

### 2.4 Analysis of the BPHCT

UPLC-MS/MS method was employed to analyse the degradation products of BPHCT. The details of the method are shown in the supplementary material.

### 2.5 Acute toxicity study in mice

The acute intraperitoneal toxicity study for calculating LD_50_ was carried out according to the OECD423 guideline (OECD(2002)) and Kaber’s method (Wang et al., 2010) with modification. The doses of BPHCT used in oral and intraperitoneal acute tests were calculated according to the original concentration of BPG solution (480 mg mL^-1^). Mice were administered by gavage with the doses of 5 mg kg^-1^, 50 mg kg^-1^, 500 mg kg^-1^, 1000 mg kg^-1^, 1500 mg kg^-1^, 2000 mg kg^-1^ and 5000 mg kg^-1^ or injected intraperitoneally with a dose level of 648.00 mg kg^-1^, 777.60 mg kg^-1^, 933.12 mg kg^-1^, 1119.74 mg kg^-1^ and 1343.69 mg kg^-1^ of BPHCT. The negative control groups were given a 0.9% normal saline solution. After 14 days observation, mortality caused by oral administration or intraperitoneal injection was calculated. The surviving mice were sacrificed for organ necropsy. To obtain the LD_50_ value and 95% confidence intervals, the following formula was used according to the Kaber’s method with modifications (Wang et al., 2010).

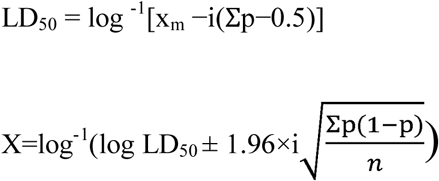

x_m_: dosage logarithm for the maximal dosage; i: the difference of dosage logarithm between two nearest groups; p: death rate of mice in each group; X: 95% confidence intervals; n: the number of the mice in each group.

### 2.6 Six months chronic toxicity study

#### 2.6.1 Experimental design

A total of 80 mice were randomly allocated into 4 groups (10 females and 10 males per group) and five mice were housed in each cage under the same conditions as described above. All mice were fed with the experimental diets every day. The weight of experimental diets provided was calculated based on the mice body weight. If the experimental diets were completely consumed before the end of a day, the normal diet would be supplied without restriction for the rest of the day. During the 6 months test period, body weight of mice was measured every 3 days in the first month and weekly during the rest of the study period.

#### 2.6.2 Clinical observation and mortality

Mice were observed twice a day for changes in fur, eyes, behaviour patterns (e.g., changes in gait or posture), morbidity and mortality, during the course of the experiment.

#### 2.6.3 Serum biochemical parameters analysis

Blood from the mice was collected in plain tubes peacefully after overnight fasting. The obtained blood was centrifuged at 3500 rpm for 10 min at 4□ and the serum was carefully transferred to a new tube and stored at −20□ for the subsequent analysis. The serum was analysed by BECKMAN COULTURE UniCelDxC 800 synchron clinical system. The clinical biochemical parameters of mice were determined including blood urea nitrogen (BUN), creatinine (CREA), total cholesterol (TC), total bilirubin (TBIL), direct bilirubin (DBIL), indirect bilirubin (IBIL), total protein (TP), albumin (ALB), globulin (GLO), alanine aminotransferase (ALT), aspartate aminotransferase (AST), alkaline phosphatase (ALP), glucose (GLU), cholinesterase (CHE), sodium (NA) and potassium (K). The contents of IgE, IL-4 and IL-13 in serum were measured by the corresponding ELISA kits.

#### 2.6.4 Histopathology

The mice were dissected to necropsy check. The organs including brain, heart, lungs, liver, spleen, kidneys, intestine and testis were carefully collected and examined macroscopically for gross lesions. The epididymides from male and femurs from female mice were also collected for the subsequent experiments. All the organs were weighted in order to calculate the ratios of organ-to-body weight and then the organs from each mouse were kept in 10% formalin for hematoxylin and eosin (HE) staining. Histological changes were examined under a light microscope.

#### 2.6.5 Sperm aberration assay

In this assay, the epididymides from male mice were cut into pieces in PBS (pH 7.2) and smears were prepared according to the method described previously (Wyrobek and Bruce, 1975). The abnormal sperms were scored and the aberration ratios were calculated.

#### 2.6.6 Polychromatic erythrocyte micronucleus formation assay

The femurs from female mice were prepared according to the method described previously (Tang et al., 2012). The polychromatic erythrocytes on the slides were observed using a light microscope. The micronucleus rates in each group were calculated.

### 2.7 Statistical analysis

Data was analysed with SPSS 20.0 program and the difference between the treatment groups and the control groups was compared. Homogeneity of variances was calculated using Levene’s test. If the variance was homogeneous, a one-way analysis of variance (ANOVA) was used to analyse the data. Otherwise, if the variance was not homogenous (p≤0.05), the identification of the statistical significance of the individual groups was tested by Dunnett’s. If the data variable is not normally distributed as required for ANOVA, Kruskal-Wallis test were performed. The statistically significance was considered to be P<0.05. Values were expressed as means±SD.

## 3. Results

### 3.1 BPHCT analysis

There were five major degradation products of BPHCT, which were benzylpenicillenic acid, N-(phenylacetyl)glycine, isobenzylpenillic acid, benzylpenillic acid and benzylpenicilloic acid. Some under baseline chromatographic peaks were observed, which might be dimer or different combination polymers of generated degradation products of BPHCT. The results are shown in the supplementary material (Supplementary Figure 1,2 and Supplementary Table 2).

### 3.2 Acute toxicity study

There was no mortality found in the oral toxicity test. Regarding observational studies, mice exhibited static, huddled up behaviour, and showed asthenia. Interestingly, mice preferred water to food in the first hour and these clinical symptoms would last for a few hours and disappear before the following day, which was correlated in a dose-dependent manner. A dose level of 1000 mg kg^-1^ was established as the no-observed-adverse-effect level (NOAEL), while the lowest-observed-adverse-effect level (LOAEL) was 1500 mg kg^-1^ for the acute oral toxicity. In the acute intraperitoneal toxicity, the symptoms of BPHCT injected mice were similar with those in the acute oral toxicity study, but some mice in the high dose group had also showed dyspnoeic symptoms or shock. The calculated LD_50_ value was 933.04 mg kg^-1^ [b.w.] and the 95% confidence intervals were 856.72-1016.15 mg kg^-1^ (Table 1).

**Table 1.** Acute toxicity of BPHCT administered through intraperitoneal injection in mice

| Dose level (mg kg<sup>-1</sup>) | Number of tested animals | Number of dead animals |
| --- | --- | --- |
| 648.00 | 10 | 0 |
| 777.60 | 10 | 2 |
| 933.12 | 10 | 5 |
| 1119.74 | 10 | 8 |
| 1343.69 | 10 | 10 |

### 3.3 Chronic toxicity study

#### 3.3.1 Effect of chronic BPHCT administration on the morbidity, mortality and body weight

Of the cohort, one female mouse and one male mouse were found crawling in circles around the cage continuously in the 60× and 600× dose group after 2 months feeding; this symptom lasted two weeks in the mouse in the 60× dose group, while it occurred in the mouse in the 600× dose group intermittently for the rest of the test. Mice in 600× dose group seemed abnormally active. At the end phase of the study, there was not a single mouse found to be dead due to drug induced toxicity.

There was a significant decrease in body weight in the female test mice compared with those in the control group (Figure 1). In the male mice, the decreased body weight was only found in the 600× dose group while increased body weight was found in the 6× and 60× dose groups compared with those in the control group.

**Figure 1.**
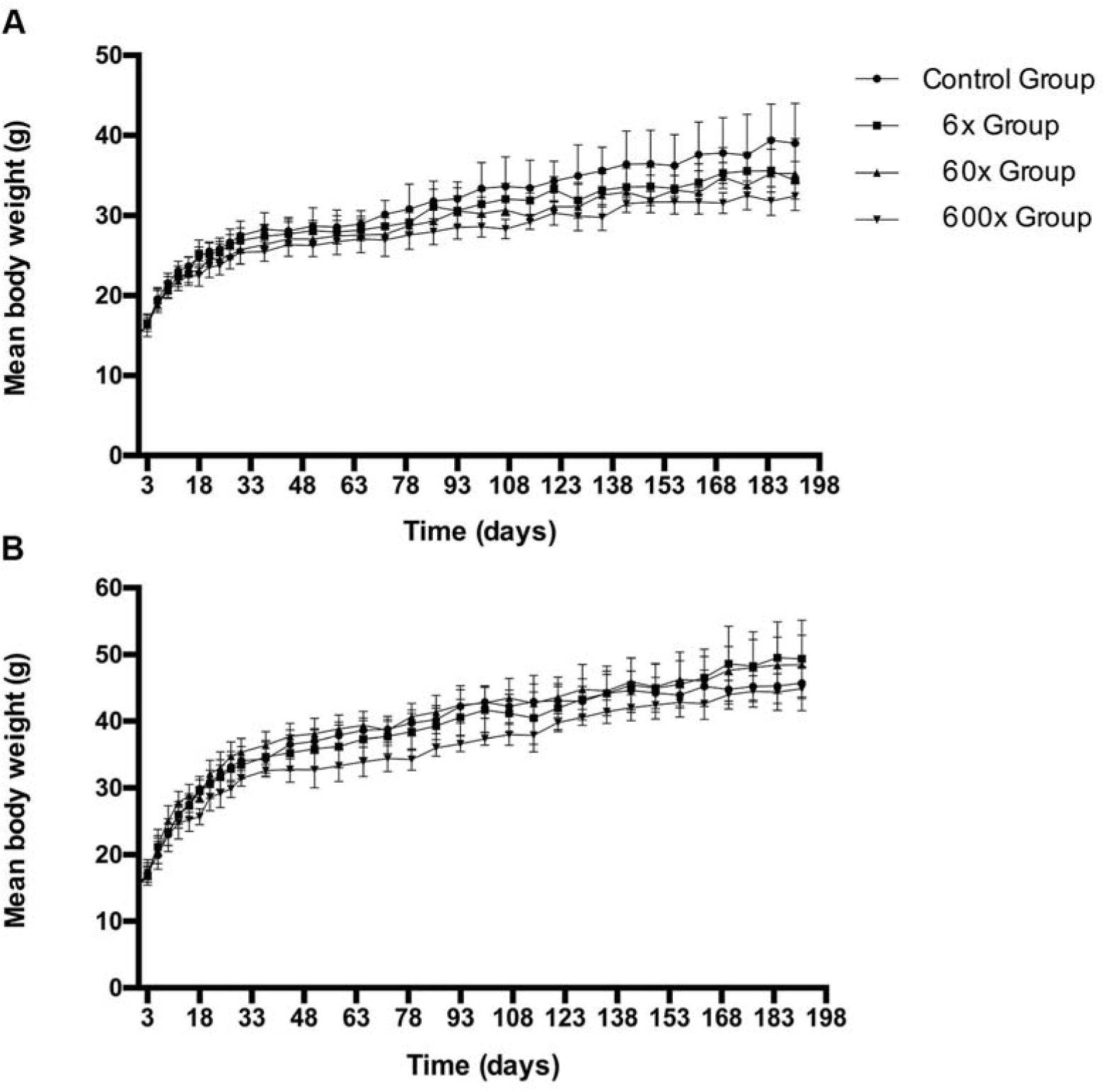
Mean body weights of mice fed with BPHCT residue in HAFP. The dose level of different groups were: control groups (0.9% saline solution), 6× groups (92.5 μg 25 g^-1^ day^-1^), 60× dose groups (925 μg 25 g^-1^ day^-1^) and 600× dose groups (9.25 mg 25 g^-1^ day^-1^) for 6 months. A. Mean body weights of female test mice. B. Mean body weights of male test mice. The data were expressed in mean±SD.

#### 3.3.2 Effect of chronic BPHCT administration on the biochemical serum parameters

The biochemical parameter profiles from control and treated mice are shown in Table 2. The level of IgE, IL-4 and 13 in both genders were raised compared with those in the control groups, but these data did not present significant difference to the data in the control groups (Table 3).

**Table 2.** Effect of 6 months oral administration of BPHCT residue in HAFP on the biochemical values of serum in mice

| Parameters | Control | 6× | 60× | 600× |
| --- | --- | --- | --- | --- |
| *Female* |  |  |  |  |
| BUN | 10.70±4.02 | 11.96±3.05 | 12.15±2.19 | 11.01±2.67 |
| CREA | 15.20±2.30 | 16.90±2.60 | 14.70±2.41 | 15.20±2.78 |
| TC | 2.38±0.56 | 2.49±0.71 | 2.12±0.75 | 2.71±0.52 |
| TBIL | 1.75±0.33 | 2.27±0.44* | 2.47±0.54** | 2.53±0.51** |
| DBIL | 0.85±0.29 | 0.93±0.31 | 0.98±0.29 | 0.97±0.40 |
| IBIL | 0.90±0.37 | 1.34±0.26** | 1.49±0.45** | 1.56±0.28** |
| TP | 56.60±3.27 | 55.60±2.55 | 56.30±2.87 | 57.20±3.58 |
| ALB | 28.40±0.97 | 27.80±1.62 | 29.30±0.95 | 29.60±0.97* |
| GLO | 27.80±2.44 | 27.80±2.04 | 26.80±1.93 | 27.60±2.88 |
| ALT | 35.60±7.57 | 35.10±7.58 | 42.70±7.17* | 48.20±6.71** |
| AST | 129.40±41.39 | 120.30±20.26 | 127.40±24.86 | 140.30±32.31 |
| ALP | 74.20±19.01 | 88.00±27.09 | 116.20±38.29** | 96.80±30.51 |
| GLU | 5.56±0.52 | 5.24±1.02 | 5.22±0.55 | 5.36±0.72 |
| CHE | 10905.10±2195.43 | 9653.40±2427.30 | 10433.10±1230.10 | 9399.20±1956.07 |
| Na | 157.85±2.03 | 156.18±2.39 | 157.97±2.53 | 156.91±1.36 |
| K | 6.67±0.63 | 7.44±0.84** | 7.27±0.51* | 6.70±0.51 |
| *Male* |  |  |  |  |
| BUN | 10.61±3.76 | 8.88±1.72 | 10.15±2.39 | 13.62±3.41* |
| CREA | 18.20±4.89 | 17.00±2.54 | 19.80±2.49 | 21.30±1.83* |
| TC | 3.49±0.72 | 3.49±0.57 | 3.78±0.58 | 3.50±0.57 |
| TBIL | 2.97±0.55 | 3.04±0.50 | 3.17±0.50 | 3.06±1.60 |
| DBIL | 1.10±0.31 | 0.90±0.36 | 1.03±0.24 | 1.17±0.54 |
| IBIL | 1.87±0.46 | 2.14±0.33 | 2.14±0.44 | 1.99±0.58 |
| TP | 52.90±4.04 | 55.70±5.85 | 56.20±4.34 | 54.20±4.49 |
| ALB | 24.90±0.99 | 26.60±1.51** | 27.10±1.10** | 26.40±0.97** |
| GLO | 27.80±2.53 | 29.10±2.51 | 29.30±1.64 | 28.10±1.20 |
| ALT | 67.00±24.72 | 44.20±5.90** | 39.10±9.26** | 42.20±8.97** |
| AST | 184.60±37.13 | 139.20±28.51** | 121.10±19.23** | 136.20±32.91** |
| ALP | 75.20±27.78 | 87.00±31.02 | 64.90±18.55 | 69.00±20.59 |
| GLU | 4.55±1.27 | 6.53±1.26** | 4.52±1.54 | 4.90±1.75 |
| CHE | 4245.50±1312.27 | 4903.40±1643.02 | 5539.00±1821.32 | 4676.30±1819.22 |
| Na | 160.03±2.42 | 158.49±2.93 | 158.98±3.02 | 159.49±2.10 |
| K | 6.76±0.77 | 7.44±0.63* | 7.55±0.74* | 7.25±0.78 |

The BPHCT with beef were offered daily to mice (n=10 per group) by feeding orally at the dose level: 6× groups, 60× groups and 600× groups, normal diets with cooked beef and 0.9% saline solution serve as the control groups. The data were expressed in mean±SD.
\*Significantly different from control group at P < 0.05. \*\*Significantly different from control group at P < 0.01.
BUN=blood urea nitrogen (mmol L<sup>-1</sup>), CREA=creatinine (umol L<sup>-1</sup>), TC=total cholesterol (mmol L<sup>-1</sup>), TBIL= total bilirubin (μmol L<sup>-1</sup>), DBIL= direct bilirubin (μmol L<sup>-1</sup>), IBIL=indirect bilirubin (μmol L<sup>-1</sup>), TP=total protein (g L<sup>-1</sup>), ALB=albumin (g L<sup>-1</sup>) GLO=globulin (g L<sup>-1</sup>), ALT=alanine aminotransferase (U L<sup>-1</sup>), AST=aspartate aminotransferase (U L<sup>-1</sup>), ALP=alkaline phosphatase (U L<sup>-1</sup>), GLU=glucose (mmol L<sup>-1</sup>), CHE=cholinesterase (U L<sup>-1</sup>), NA=sodium (mmol L<sup>-1</sup>), K=potassium (mmol L<sup>-1</sup>).

**Table 3.** Effect of 6 months oral administration of BPHCT residue in HAFP on the contents of Immunoglobulin E and Interleukins in the serum of mice

| Dose level |  |  | Control | 6× | 60× | 600× |
| --- | --- | --- | --- | --- | --- | --- |
| Parameters | IgE (μg ml<sup>-1</sup>) | Male | 4.50±2.05 | 3.82±3.99 | 5.43±4.81 | 5.36±3.21 |
|  |  | Female | 5.24±6.33 | 4.78±5.59 | 4.51±2.03 | 6.17±5.30 |
|  | IL-4 (pg ml<sup>-1</sup>) | Male | 22.56±4.55 | 23.60±11.37 | 26.63±5.11 | 25.93±5.53 |
|  |  | Female | 17.85±1.49 | 18.93±2.82 | 18.63±0.87 | 19.16±2.37 |
|  | IL-13 (pg ml<sup>-1</sup>) | Male | 20.54±1.47 | 19.66±0.53 | 21.02±1.73 | 28.60±13.04 |
|  |  | Female | 21.32±3.34 | 26.77±12.32 | 25.43±9.04 | 25.09±13.22 |

Data were expressed in mean±SD, n=7.
\*Significantly different from control group at P < 0.05. \*\*Significantly different from control group at P < 0.01.

#### 3.3.3 Effect of chronic BPHCT administration on Organ ratios and Histopathology

The data of organ ratios from control and treated mice are shown in the Table 4. The brain ratios in male and female test mice were increased but were not statistically different.

**Table 4.** Effect of 6 months oral administration of BPHCT residue in HAFP on the organ to body weight ratio in mice

| Dose level |  |  | Control | 6× | 60× | 600× |
| --- | --- | --- | --- | --- | --- | --- |
| Brain | Male |  | 11.00±1.13 | 10.39±2.14 | 11.32±1.42 | 11.36±1.69 |
|  | Female |  | 13.01±1.27 | 14.61±1.42 | 14.56±2.83 | 14.96±1.09 |
| Heart | Male |  | 7.16±2.11 | 6.72±1.06 | 6.95±1.14 | 6.27±1.00 |
|  | Female |  | 6.51±1.69 | 6.58±1.25 | 6.14±1.18 | 6.82±1.14 |
| Liver | Male |  | 45.64±5.81 | 40.69±7.87 | 38.85±2.67** | 41.36±4.56 |
|  | Female |  | 44.76±11.10 | 46.49±4.73 | 41.44±4.60 | 45.63±3.83 |
| Spleen | Male |  | 2.98±0.52 | 2.52±0.60 | 2.61±0.76 | 3.29±1.18 |
|  | Female |  | 3.07±0.79 | 3.58±0.54 | 3.61±0.44* | 3.91±0.49** |
| Lungs | Male |  | 7.15±1.11 | 6.99±0.48 | 6.44±1.06 | 6.66±0.88 |
|  | Female |  | 7.24±1.13 | 8.00±1.05 | 7.18±1.14 | 7.31±0.72 |
| Kidneys | Male |  | 16.46±2.27 | 16.26±1.64 | 15.79±2.02 | 16.05±2.25 |
|  | Female |  | 12.34±1.85 | 13.58±1.37 | 13.75±1.74 | 13.28±1.38 |
| Intestine | Male |  | 76.48±7.56 | 76.76±5.86 | 80.66±6.61 | 87.09±16.25* |
|  | Female |  | 101.69±11.36 | 100.49±12.97 | 106.35±16.38 | 118.02±10.36** |
| Testes | Male |  | 7.11±1.19 | 7.14±1.35 | 7.33±2.31 | 6.66±1.67 |

Data were showed with means±SD. The organ ratios were calculated using organ weight/body weight × 1000, n=10.
\*Significantly different from control group at P< 0.05. \*\*Significantly different from control group at P< 0.01.

In the 600× dose group, diffusely hepatocytes showed eosinophilic granular cytoplasm, individual hepatocyte, necrosis, lymphocytic and histiocytic inflammatory infiltrate in the liver parenchyma (Figure 2). The pulmonary changes consisted of multifocal lymphocytic, histiocytic and plasmacytic inflammatory infiltration in the parenchyma (Figure 3). Furthermore, in the testis, seminiferous tubules were absent or fewer cellular than the control cases, while the interstitium was broadened. There was diffuse severe necrosis of the spermatogenic lineage cells, decreased number of sustentacular cells and spermatids were absent or infrequent (Figure 4). The changes mentioned above were found in 30% percentage of mice in 600× dose group. Histopathological changes were not found in the control and 6× dose groups and few in the 60× dose groups after 6 months treatment.

**Figure 2.**
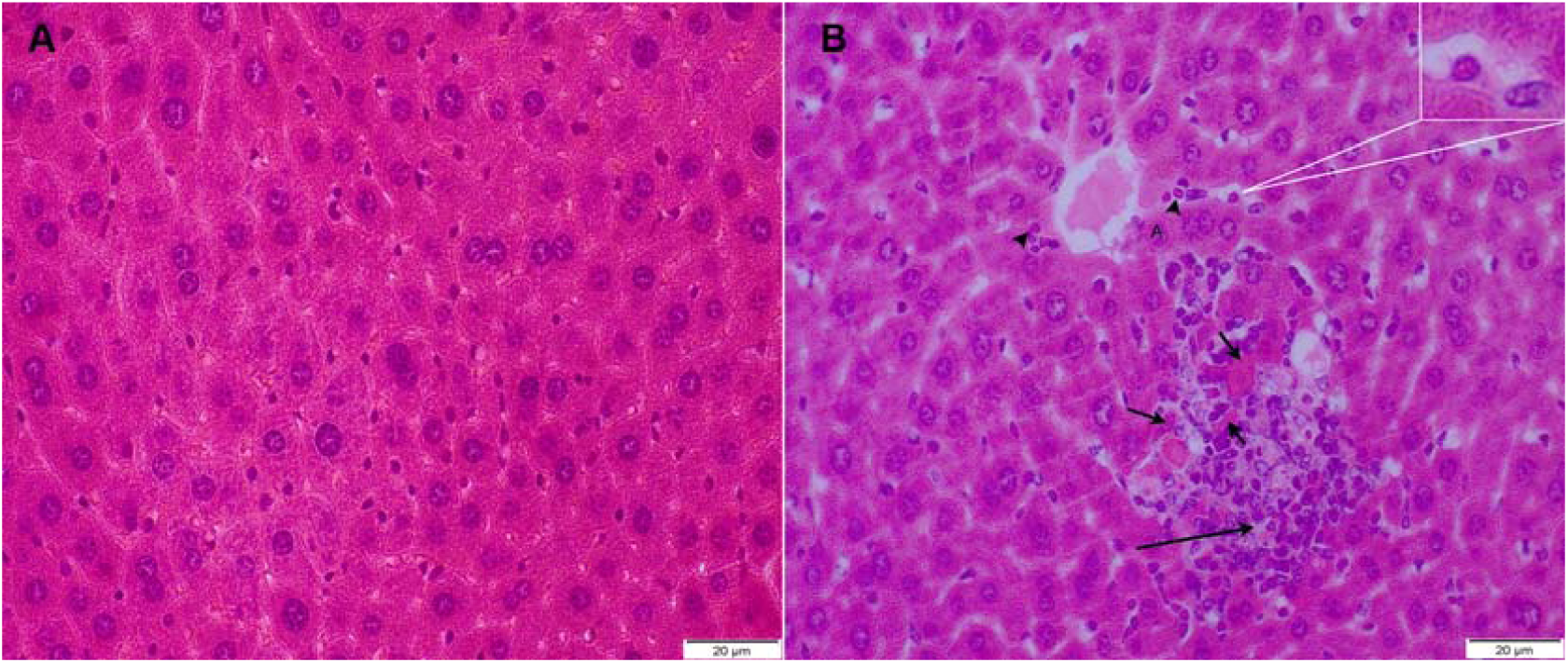
Histopathological changes in the liver tissue from control group (A) and 600× dose group (B) (× 400). The lymphocytic and histiocytic inflammatory infiltration (long arrow), necrosis hepatocyte (short arrows) and eosinophilic granular cytoplasm (arrow head and the magnified box) were observed in the liver parenchyma in the 600× dose group. No characteristic histopathological changes were observed in the liver of the control mice.

**Figure 3.**
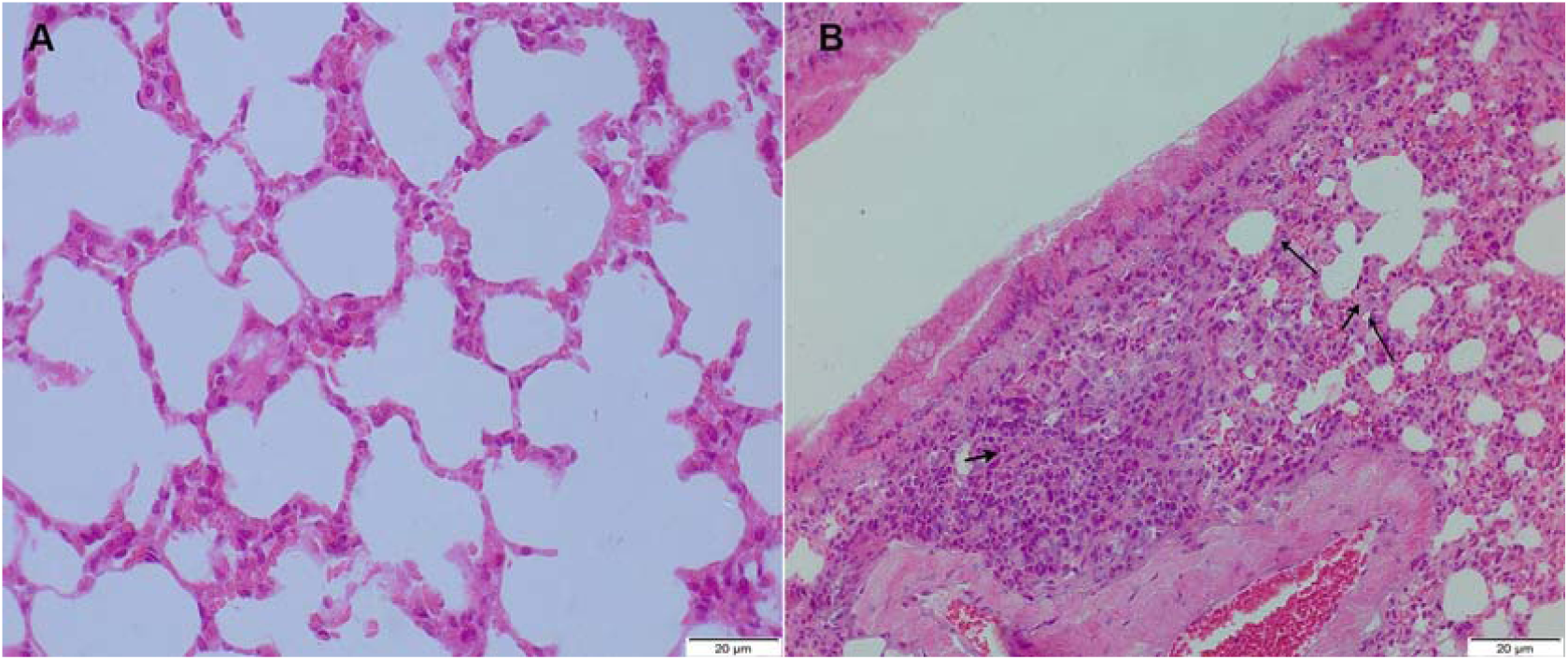
Histopathological changes in the pulmonary tissue from control group (A) and 600× dose group (B) (× 400). Multifocal lymphocytic (long arrow), histiocytic and plasmacytic inflammatory infiltration (short arrows) were observed in the parenchyma in the 600× dose group. No characteristic histopathological changes were observed in pulmonary tissue of the control mice.

**Figure 4.**
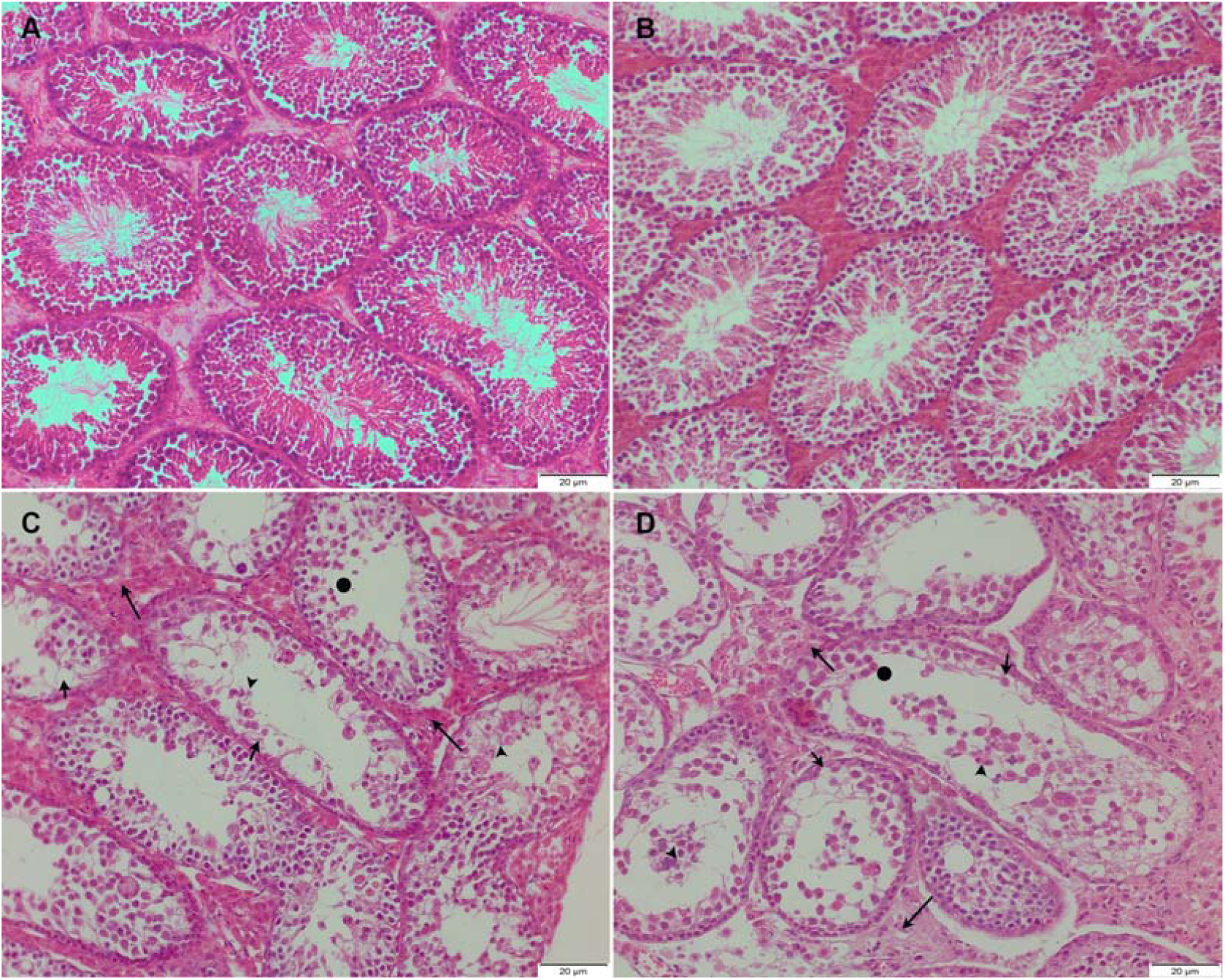
Histopathological changes in the testis tissue from control group (A), 6× group (B), 60× dose group (C) and 600× dose group (D) (× 400). In the 600× dose groups, seminiferous tubules (long arrows) were loosed and interstitial broadening. The decreased population of sustentacular cells (short arrows), necrotic spermatids (arrow head) and the fewer spermatids remained within the lobules (the circles) were observed. While, these histopathlogical changes could be observed less in the 60× dose group. No characteristic histopathological changes were observed in testis of Control and 6× groups.

#### 3.3.4 Effect of chronic BPHCT administration on sperm aberration rate

Several sperm abnormalities including hookless and amorphous were observed with a high frequency in the 60× and 600× dose groups, and at a lower frequency in the 6× dose group administered with BPHCT residue in HAFP orally for 6 months (Figure 5, Table 5).

**Table 5.** Effect of 6 months oral administration of BPHCT residue in HAFP on aberration rates of sperm in male mice and polychromatic erythrocyte micronucleus formation rates in female mice

| Dose level | Frequency of aberration (%) | MNPCE (%) |
| --- | --- | --- |
| Control | 1.68±0.22 | 8.69±1.19 |
| 6× | 2.11±0.65 | 10.52±1.33 |
| 60× | 9.35±1.30** | 10.23±4.08 |
| 600× | 19.41±3.28** | 15.97±1.13** |

Data were expressed in mean±SD, 5000 cells/group, n=5, MNPCE= micronucleated polychromatic erythrocytes.
\*Significantly different from control group at P < 0.05. \*\*Significantly different from control group at P < 0.01.

**Figure 5.**
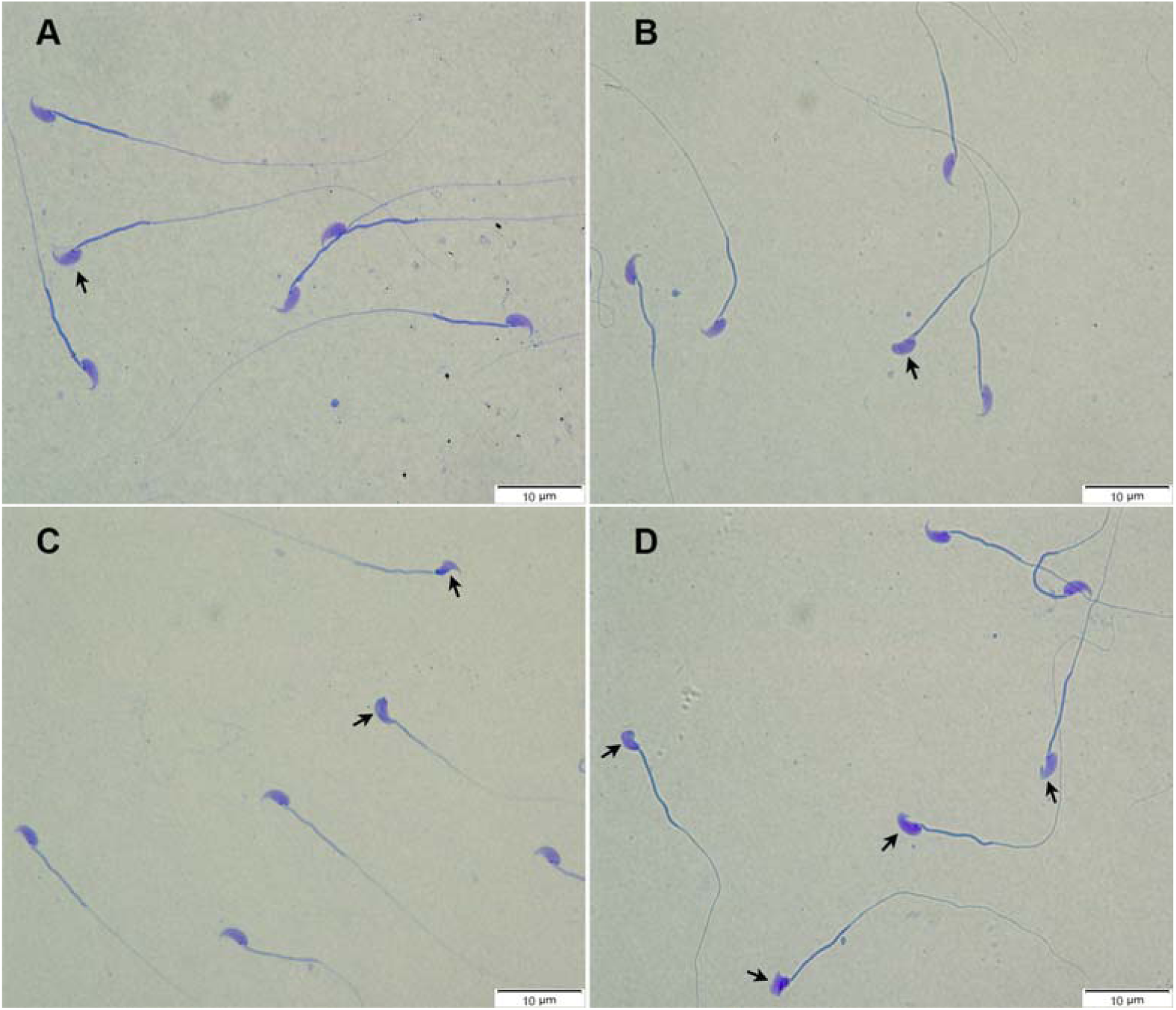
Selected microphotographs of sperms in male mice after 6 months oral administration of BPHCT residue in HAFP (×1000). Compared with the control (A) and 6× (B) groups, a significant increase in the sperm aberration rate, such as hookless or amorphous acrosome (short arrows), were observed in the 60× (C) and 600× (D) dose groups.

#### 3.3.5 Effect of chronic BPHCT administration on Polychromatic erythrocyte micronucleus formation

Compared with the control group, no statistically significant increase in the micronucleus formation was observed in mice feed with experimental diets in 6× and 60× dose of BPHCT for 6 months (Table 5). However, the rate of micronucleus formation was readily increased in the 600× dose BPHCT group compared with those in the control group (Figure 6).

**Figure 6.**
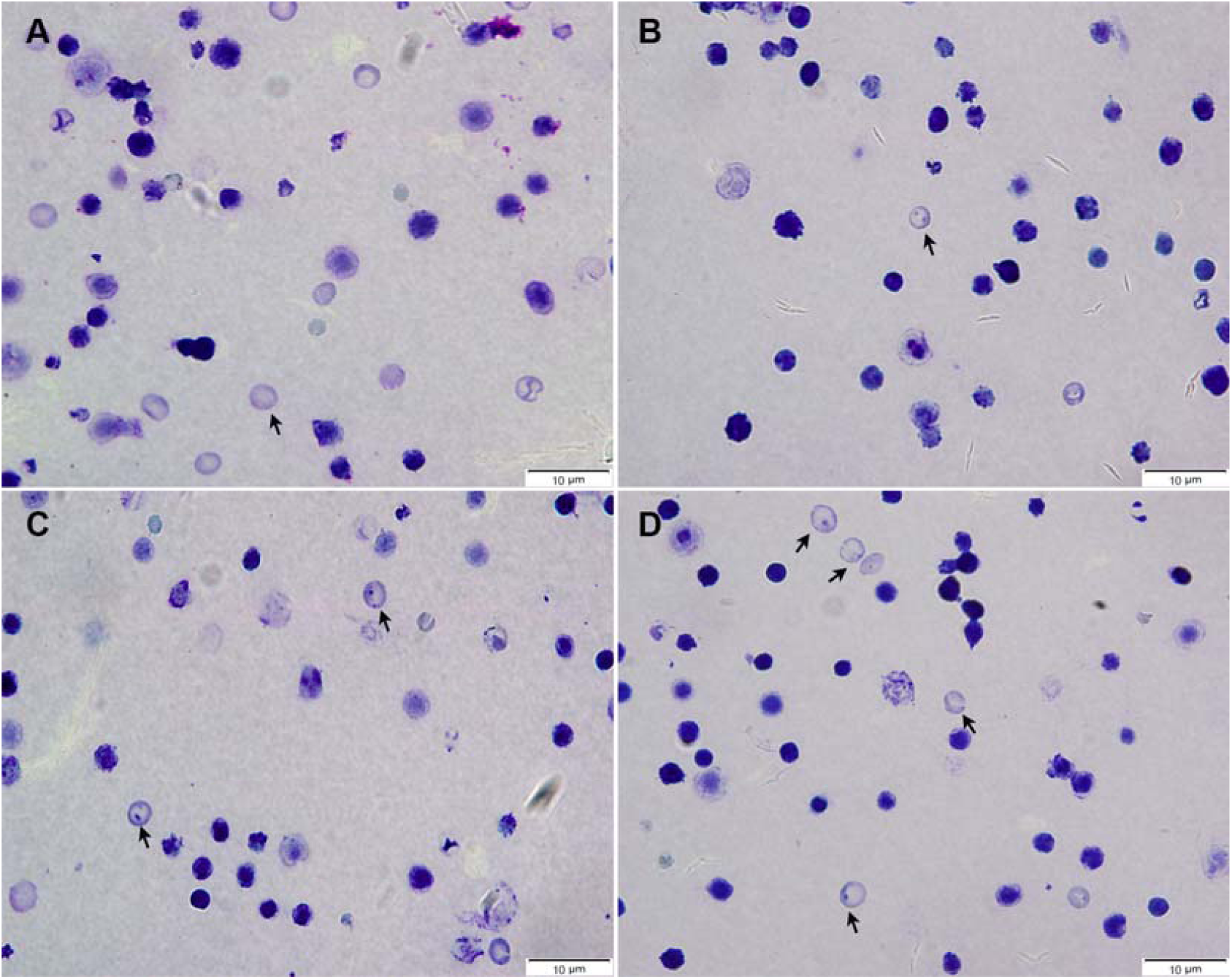
Selected microphotographs of polychromatic erythrocyte micronucleus in female mice after 6 months oral administration of BPHCT residue in HAFP (×1000). A (Control group) presents few polychromatic erythrocyte micronucleus, short arrow shows the normal polychromatic erythrocytes in the picture. B (6× group) and C (60× dose group) present a lower rate of polychromatic erythrocyte micronucleus (short arrows) formation. A significant increase in the micronucleus formation was observed in D (600× dose group). One or two micronucleus could be observed in one cell.

## 4. Discussion

Recent studies have showed that animal derived products containing penicillin residues exceeding the regulatory safety level have caused an increasing health concern to consumers globally (Baynes et al., 2016). Many studies have demonstrated that the structure of penicillin is unstable at high temperatures (Tian et al., 2017). However, very few studies have reported the toxicity of penicillin heated to cooking temperature, and the safety of penicillin residues in HAFP. Hence, assessment of the toxicity of penicillin residues in HAFP is required and necessary to advise protocols pertaining to public health. In this study, the toxic effects of BPHCT residue were investigated in a mouse model.

In the acute toxicity study, the LD_50_ of BPHCT was 933.04 mg kg^-1^. A previous study reported that the LD_50_ of BPG was 3500 mg kg^-1^ injected intraperitoneally (Hobby, 1968). The result suggested that the toxicity of BPHCT was 3.75 times higher than its prototype.

It has been reported that a decrease in body weight has been used to demonstrate the adverse effects of chemicals and drugs (Teo et al., 2002). In the chronic toxicity study, we suggest that the decreases in the mean body weights in test mice were the result of the adverse effects caused by BPHCT. However, the reason for the promotion of growth in 60× dose male group was not clear, which should be subject to further investigation. Abnormal activity and crawling in circles are likely to be neurological symptoms, which may lead to the suppression of body weight growth in the test animals. The BPHCT residues in HAFP may possibly be responsible for the decreased growth and abnormal symptoms in the test mice.

Haematological analysis is a relevant part of toxicity evaluation for its higher predictive value for human toxicity (Olson et al., 2000). The levels of BUN and CREA in serum are important markers for kidney damage. The level of BUN increases when the blood volume and Glomerular Filtration Rate decrease (Lee et al., 2007). The level of CREA increases as the filtration rate decreases (Wang et al., 2015a). Present results showed that the levels of BUN and CREA in the serum of 600× dose male group but not the female group were significantly increased compared with those in the control group. High levels of BUN and CREA indicated that BPHCT in HAFP might affect the Glomerular Filtration function and even cause the kidney damage or necrosis to male kidney cells.

The two transaminases AST and ALT are used as biomarkers to evaluate the function of liver and their elevated values in the serum are indicative of liver damage (Tennekoon et al., 1991). Our results showed increased levels of ALT, AST in female test mice compared with those in the control group, indicating liver damage in these animals. Apart from ALT and AST, the other biomarker-TBIL, is also used to evaluate liver function. TBIL is consisted of DBIL and IBIL, and a high bilirubin concentration may indicate the liver dysfunction (Wang et al., 2014). In our study, these serum parameters changed markedly in test mice compared with those in control groups, which indicated that the liver function might be affected in BPHCT treated mice. Taken together, the results suggested that high dose of BPHCT residues might be toxic to liver, affecting liver function.

Histopathological assessment of the liver showed that the hepatocyte granular cytoplasm, necrotic changes and hepatitis were found in liver tissues in mice in the 600× dose cohort. These histopathological findings may have a correlation with changes in the level of ALT, AST and the bilirubin in the serum of 600× dose mice. These evidences may prove the potential hepatotoxicity of BPHCT. According to our UPLC-MS/MS results, BPG broke down to form benzylpenicillenic acid, N-(phenylacetyl)glycine, isobenzylpenillic acid, benzylpenillic acid, benzylpenicilloic acid and polymers that similar to the result reported previously (Depaolis et al., 1977). These degradation products are highly immunogenic agents due to the rupture of β-lactam ring and the penicilloyl group formation, which bind with protein via a covalent bond to form a completed antigen (Levine and Price, 1964; Wal et al., 1975). We theorised that the inflammatory cell infiltration and consolidation changes in lungs might be the result of increasing inflammatory cell stimulated by penicillin degradation products binding proteins. According to our results, parameters of IgE and interleukins in the test groups increased with different degrees but not significantly. Here, we hypothesise that the presence of relatively high level of IgE and interleukins in the serum could be specific for the degradation products of BPHCT. Interestingly, we didn’t find obvious histopathological evidence in the brain to explain the neurological symptoms of test mice. However, it had been reported that penicillin was neurotoxic (Nicholls, 1980). The observed neurological signs of test mice should not be ignored and the mechanism needs further elucidation.

We know that naturally occurring mistakes or exogenous factors such as genotoxicity chemicals and irradiation may cause sperm aberration or spermatogenic dysfunction during the differentiation (Bruce et al., 1974). In our test, we observed that the sperm aberration rates in the 60× and 600× feeding groups were significantly increased compared with those of the control group. The two test groups displayed characteristic shapes indicating morphological changes, including hookless and amorphous heads. This suggests that high concentration of BPHCT residue might affect spermatogenic tissues. The histopathological study of the testis also showed dead spermatogonia cells and the decreased population of sertoli and sperm cells in spermatogenic tissues, which are the evidence to support the possible toxicity of BPHCT on spermatogenic tissues in mice. On the other hand, the micronucleus assay is considered a preferred method for assessment of genotoxic effect and chromosome damage caused by exposure to ionizing radiation or carcinogenic chemicals (Chauhan et al., 2000; Fenech, 2000). Multiple micronuclei may form as the chromosome lagged behind and fail to incorporate into daughter nuclei during the anaphase of mitosis. In the polychromatic erythrocyte micronucleus assay, the formation rate of micronucleus in the bone marrow cells from 600× dose group increased significantly compared with that of the control group. Based on our results, we suggest that higher dose of BPHCT residue might interfere a lagging acentric chromosome fragment and be a potential genotoxicity agent. Therefore, our results suggested that sperm aberration and micronucleus formation caused by degradation products of BPHCT in mice indicates possibilities of sperm toxicity and genotoxicity.

This work has clearly demonstrated potential toxic effects of BPHCT *in vivo*. We investigated the potential toxicity of HAFP contaminated with penicillin residues, which has not been studied and remained as a blind spot of the public health authorities and consumers. The five major thermal degradation products of BPHCT contain different functional groups such as carboxyl-, hydroxyl-, sulfhydryl- and carbonyl-group etc., which could explain the observed toxicity in mice. Polymers formed in the process also need to be paid more attention to due to their complex structures. Among these products, only the toxicity of benzylpenicilloic acid has been evaluated *in vivo* and *in vitro* (Cui et al., 2018). More studies are needed to address the toxic effects of these degradation products of BPHCT.

According to our pilot study, we found that low dose of BPHCT doesn’t show toxic effects in mice. The high doses of BPG (60× and 600× dose) were used in the chronic study to evaluate the potential toxicity after heat treatment but not death or severe suffering based on the OECD guidelines, even though the concentration is significantly higher than those observed in the environment. In some countries, especially in developing countries and some rural areas, BPG residues are a major issue due to its illegal use and poor management of withdrawal period (Babapour et al., 2012). Therefore, the possibility of exposing high dose of BPG residues to the public should not be ignored.

In another aspect, there are huge differences in the policies of penicillin usage and residues levels worldwide. Present results may be not valuable to those countries that do not allow the use of penicillin in animal husbandry. Nevertheless, this study systematically reported the toxicity of penicillin residues in animal derived products heated to cooking temperature and their potential threats to public health. It also suggested that although rarely reported, the potential harm posed by other antibiotic residues in food should be investigated.

## 5. Conclusions

In summary, the toxicity of BPG is increased by over 3.75 times as a result of heat-treatment. The BPG has no significant side effect at the 6× dose level in edible animal tissue after heat-treatment. However, over 60× or 600× dose level may lead to various toxicities, particularly the potential hepatotoxicity and pulmonary toxicity, as well as sperm aberration and micronucleus formation after long time exposure. Taking the thermal process into account, this study provides the toxicity evaluation of antibiotics residue in animal derived products, which would inform the legislation on antibiotic usage guidelines. It is necessary for the government to perform a strict supervision and withdrawal period on the use of penicillin and other heat-unstable antibiotics in animal husbandry.

## Acknowledgements

We thank Dr. Mark. A. Holmes (University of Cambridge, Dept of Veterinary Medicine, Madingley Road, Cambridge CB3 0ES, UK) for kindly revising the manuscript. This work was supported by the National Natural Science Foundation of Jilin Province, P.R.China (grant no. 201701035JC) and Key Projects of Jilin Province Science and Technology Development Plan (20140204065NY). Cheng Cui was supported by the China Scholarship Council.

